# Distributed Compact Plasma Reactor Sterilization for Planetary Protection in Space Missions

**DOI:** 10.1101/2022.11.14.516453

**Authors:** Bhaswati Choudhury, Tamara Revazishvili, Maria Lozada, Sarthak Roy, Emma Noelle Mastro, Sherlie Portugal, Subrata Roy

**Affiliations:** SurfPlasma, Inc, 32601, Gainesville, USA; Emerging Pathogens Institute, University of Florida, Gainesville, Fl-32611, USA; School of Electrical Engineering, Technological University of Panama, Panama City, Panama; Dept. of Mechanical and Aerospace Engineering, University of Florida, Gainesville, Fl-32611, USA

**Keywords:** plasma, planetary protection, ozone, flow actuation, sterilization, dielectric barrier discharge

## Abstract

This paper presents a proof-of-concept study establishing effectiveness of the Active Plasma Sterilizer (APS) for sterilization in planetary protection. The APS uses Compact Portable Plasma Reactors (CPPRs) to produce surface dielectric barrier discharge, a type of cold plasma, using ambient air to generate and distribute reactive species like ozone used for decontamination. Sterilization tests were performed with pathogenic bacteria (Escherichia coli and Bacillus subtilis) on materials (Aluminum, Polycarbonate, Kevlar and Orthofabric) relevant to space missions. Results show that the APS can achieve 4 to 5 log reductions of pathogenic bacteria on four selected materials, simultaneously at 11 points within 30 minutes, using power of 13.2 ± 2.22 W. Spatial sterilization data shows the APS can uniformly sterilize several areas of a contaminated surface within 30 minutes. Ozone penetration through Kevlar and Orthofabric layers was achieved using the CPPR with no external agent assisting penetration. Preliminary material compatibility tests with SEM analysis of the APS exposed materials showed no significant material damage. Thus, this study shows the potential of the APS as a light-weight sustainable sterilization technology for planetary protection with advantages of uniform spatial decontamination, low processing temperatures, low exposure times, material compatibility and the ability to disinfect porous surfaces.

## Introduction

With the advancement of space exploration comes the need to develop technologies and practices that protect the integrity of scientific investigations in space. Planetary protection is the practice of preventing cross contamination between earth and any celestial body of interest in a mission. This ensures credibility of the data collected during space missions while protecting the earth from extra-terrestrial organisms (backward contamination) and vice versa (forward contamination) [1].

Although contamination control technologies for planetary protection are established for uncrewed missions, scientists are exploring technologies for crewed missions where microbial mitigation have increased complexity levels due to risk of recontamination by human contact [1,2]. Furthermore, existing technologies approved by National Aeronautics and Space Administration (NASA) and the European Cooperation for Space Standardization (ECSS) for uncrewed missions-dry heat microbial reduction (DHMR) and vaporized hydrogen peroxide (VHP) bio decontamination, have certain drawbacks [3–7]. The ECSS reported that DHMR can damage heat sensitive materials while VHP can cause detrimental material alteration [6,7]. Knowledge gaps related to required concentrations in VHP, it’s delivery mechanism and material compatibility have also hindered its utilization in space missions [2]. Thus, there is a need for alternative technologies that can overcome disadvantages of currently approved methods for uncrewed missions while addressing microbial mitigation in crewed missions.

Researchers have identified non-thermal plasma (NTP) as a potential alternative to traditional decontamination methods used in planetary protection and other areas like food preservation and surface sterilization [8–11]. Dielectric barrier discharge (DBD) is a type of NTP which has been proven effective against a variety of microbes including bacterial endospores which is crucial for planetary protection applications [9–14]. DBD is the electrical discharge formed between two electrodes separated by a dielectric barrier when a high enough AC voltage is applied across them. As the name suggests, surface DBD (SDBD) is the DBD formed on a surface. SDBD is unique as it causes generation of reactive species while influencing the flow of the surrounding gas without employing any moving parts. DBD decontamination occurs through direct contact with discharge or indirect contact with reactive species formed by the discharge [15]. Indirect DBD treatment is advantageous for treating hidden surfaces and surfaces relatively larger than the discharge area. When SDBD is generated in air, reactive oxygen and nitrogen species (RONS) are formed that can destroy microorganisms like bacteria, viruses and fungi. One such species is ozone which is known to cause microbial inactivation and is the longest living species in comparison to other RONS formed during SDBD generation in air [16,17]. The mechanism of such microbial inactivation occurs through a progressive set of complex reactions that lead to destruction of cellular surface, leakage of cellular surface, leakage of cellular components and cell lysis, finally inactivating the micro-organism [18].

This paper discusses the proof-of-concept study performed under a NASA contract to establish the effectiveness of the Active Plasma Sterilizer (APS) - a compact and energy efficient sterilization system with inbuilt ozone generation and mixing capability and evaluates it for sterilization applications pertaining to planetary protection [19]. The Active Plasma Sterilizer (APS) consists of dry, versatile, reusable and modularly scalable Compact Portable Plasma Reactors (CPPRs) which produce SDBD using a few watts of electrical power across electrodes separated by a dielectric medium [20,21]. It uses the concept of achieving spatially distributed decontamination using a synergistic combination of SDBD ozone generation and flow actuation for distributing and mixing generated ozone without using external mixing agents [15, 22]. Doing so maximizes utilization of ozone generated for rapid decontamination while reducing ozone requirements and residual concentrations [15]. To the best of our knowledge, APS is the first device designed to achieve spatially distributed decontamination using SDBD reactors. This study evaluates the first APS prototype – APS.V0 by determining its sterilization efficacy, ozone CT (concentration x time) requirements, power requirements, penetration capability through desired materials and material compatibility. Sterilization tests were performed with bacteria (*Escherichia coli and Bacillus subtilis*) and materials (Aluminum, Polycarbonate, Kevlar and Orthofabric) relevant to space missions [1,23,24]. Results show that the APS prototype can achieve complete killing (4 to 5 log reductions) of pathogenic bacteria (*Escherichia coli* and *Bacillus subtilis*) on 4 selected materials simultaneously at 11 points inside the chamber within 30 minutes using 13.2 W total power consumption. Additionally, successful ozone penetration for single and combined fabric layers was established without using an external agent. Further, preliminary material compatibility tests with SEM analysis of 4 selected materials exposed to ozone CT values required for sterilization in the APS showed no significant material damage.

The APS is a lightweight, low-cost, non-thermal and rapid decontamination solution for potential integration into spacecraft or platform subsystems. It aims to address the disadvantages related to currently approved methods for space missions like high processing temperatures and material incompatibility. Additionally, the APS may be used to achieve microbial mitigation in crewed missions. In comparison to large and high-power consuming DBD ozone generation systems that already exist, the APS is unique with its (a) compact, lightweight and low energy power supply and (b) SDBD flow actuation aided ozone mixing leading to rapid decontamination with lower ozone requirements and lower residual ozone [15,20]. Potential applications include sterilization of spacecraft components and subsystems, ground-based contamination control that can withstand testing operations and in-flight component cross-contamination control. Some application examples are sterilization of sample acquisition equipment (drills, etc.), surfaces and space suits pre- and post-launch.

## Materials and Methods

### Design of the APS prototype (APS.V0)

#### Sterilization box

A custom polycarbonate box with internal dimensions 1 ft long × 1 ft wide × 1 ft high and wall thickness of 0.0065 ft was built. A layer of foam insulation adhesive was placed between the lid and sidewalls of the box to seal it from the external environment. Polycarbonate was chosen as it is non-reactive to ozone. For measuring decontamination achieved at internal points of the chamber, holes were drilled in the sidewalls to facilitate suspension of inoculated coupons inside the box as shown in Figure 1. Three planes were selected to represent volume occupied by an object placed in the box: Plane 1 (P1), Plane 2 (P2) and Plane 3 (P3) positioned at d1= 8 cm, d2=11.5 cm and d3= 16 cm from the bottom of the box, respectively. The (P, Q, R) coordinate system is used to explain the decontamination measurement grid a later section. The drilled holes also provided access to a measurement probe for ozone data collection inside the box.

**Figure 1.**
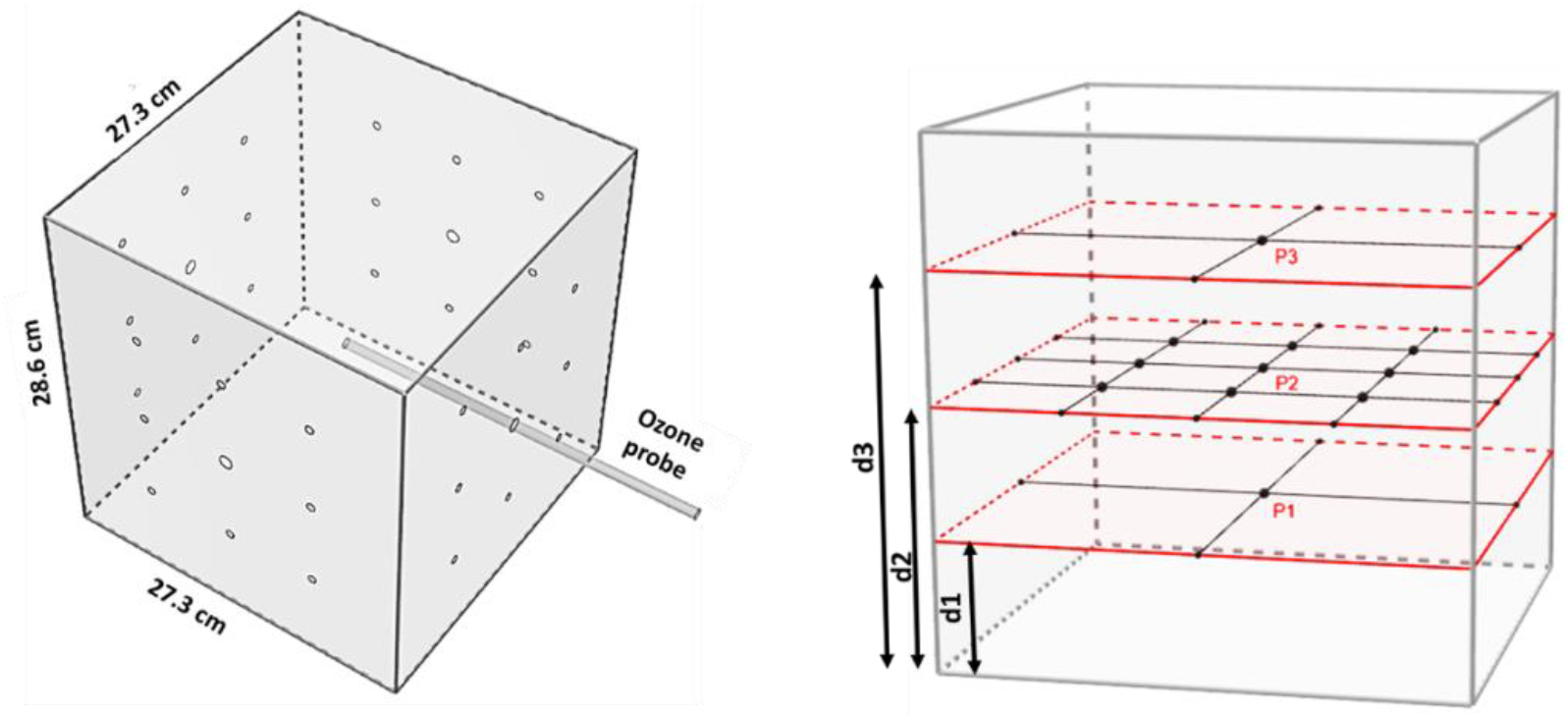
Schematic of APS sterilization box showing drilled holes for accessing decontamination measurement locations in three planes inside the box. This set up also provided access to the center of the box for ozone data collection using a probe connected to a ozone monitor.

#### Compact Portable Plasma Reactor (CPPR)

The CPPRs incorporated into the APS sterilization box were designed to be compact and portable [20]. Two components of the CPPR are: (a) the reactor panel consisting of two sets of copper electrodes separated by a dielectric medium and (b) the compact power supply circuit which converts low DC voltage to high AC voltage required for creating dielectric barrier discharge on the reactor panel surface. The copper electrode sets were 35 μm thick separated by a dielectric material: hydrocarbon/ceramic (RO4350B™[25]) composite - 0.76 mm thick with 3.48 dielectric constant. The CPPR power supply, also known as the Active plasma module, is a compact circuit module with a volume of 48 cubic centimeters and weight of 55 grams [21]. It runs on 25V DC power supply and consumes an average of 2.2+0.37 Watts power. Details of the power measurements of this active plasma module can be found in a previously published study [21]. The power supplies were encased in electrically insulating cases for added safety. The CPPR reactor panel designs were selected based on their flow actuation capabilities responsible for distributing the ozone generated by them [15,22]. Two reactor panel designs with contrasting flow actuation capabilities were used for better distribution of generated ozone in the box - the comb reactor and the fan reactor. The two panel designs are shown in Figure 2. The comb reactor panel results in a 2d flow distribution with a dominant wall jet in the direction from the shaft towards the teeth tips while the Fan reactor results in a 3d flow distribution forming a swirl flow spreading vertically upwards and outwards from the center of the panel. Details of the flow actuation and ozone distribution caused by the reactor panel designs can be found in a recently published paper on Smart DBD plasma decontamination [15]. Geometric details of the reactor panels are kept same as those used for ozone studies in the development of the SDBD Fan configuration [22].

**Figure 2.**
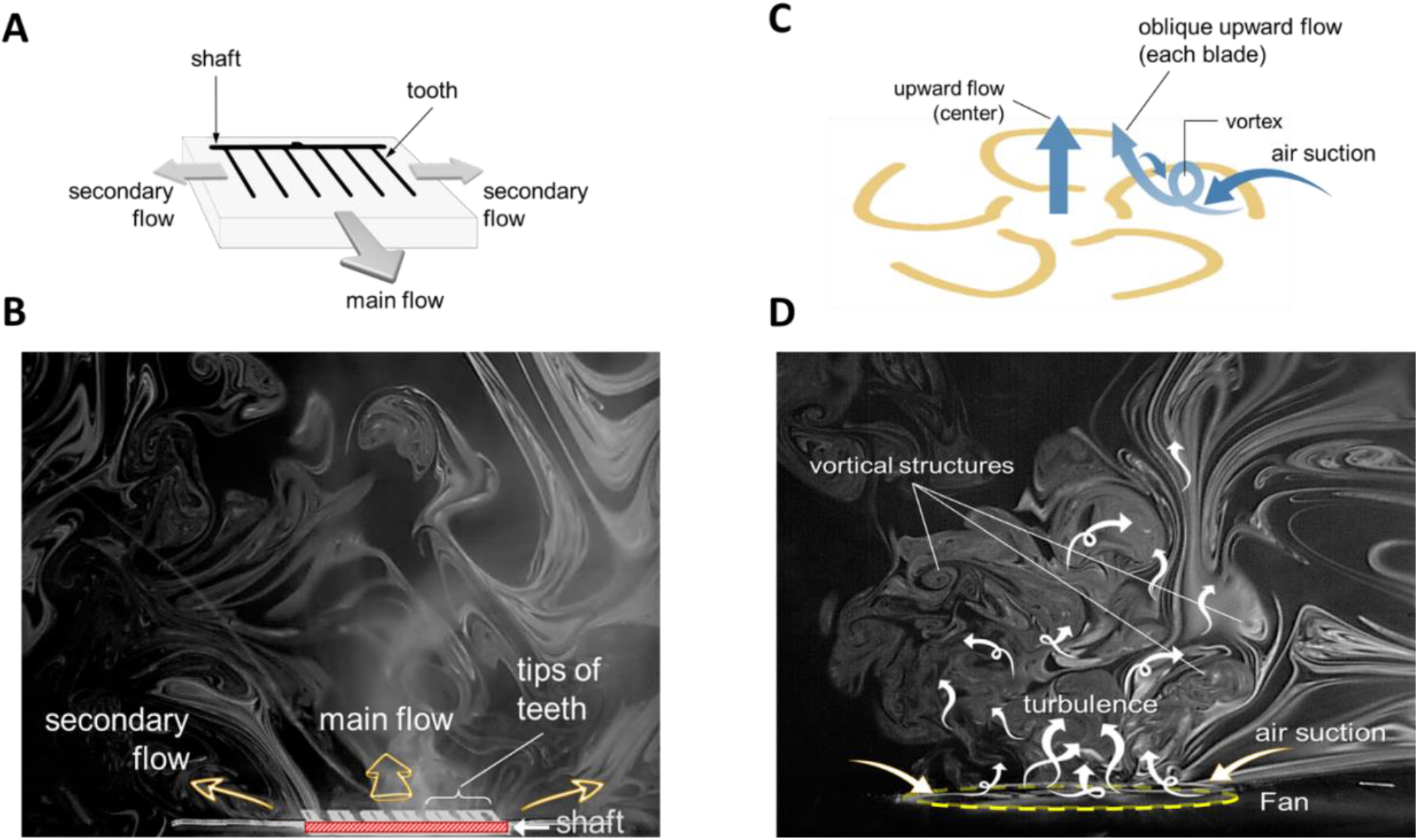
Smoke flow visualizations showing plasma reactor flow distribution with two reactor designs: Comb and Fan. (a) Comb reactor flow actuation showing the direction of three wall jets with the main wall jet flow from the shaft towards the teeth tips, (b) smoke flow visualization of Comb reactor actuated flow, (c) Fan reactor flow actuation by the fan blades which interacts to result in overall conical flow thrusted upwards from the reactor base, (d) smoke flow visualization of Fan reactor actuated flow. Figure adapted from previous publication [15].

#### Integration of the CPPRs into the sterilization box

4 to 6 CPPRs were integrated in the sterilization box as shown in Figure 3. The placement and orientation of the reactors was based on the expected ozone distribution each CPPR reactor panel produces [15]. Three CPPRs with Fan reactor panels were placed on the lid in an equilateral triangle placement to achieve the effect of vortical showers of ozone on the internal volume of the chamber. One CPPR with the Comb reactor panel was placed at the bottom of the chamber with a 2 cm offset from the center. The Comb reactor panel was placed such that the dominant wall jet from the shaft towards the teeth tips was directed upwards [15]. This ensured ozone transportation from the bottom to the top which is expected to diffuse through the internal chamber volume. The offset was given to avoid blocking of the wall jet by the coupons placed at the center points in planes 1,2 and 3. The integration of the CPPRs in the sterilization chamber was done such that the number of CPPRs powered up during the experiments could be controlled.

**Figure 3.**
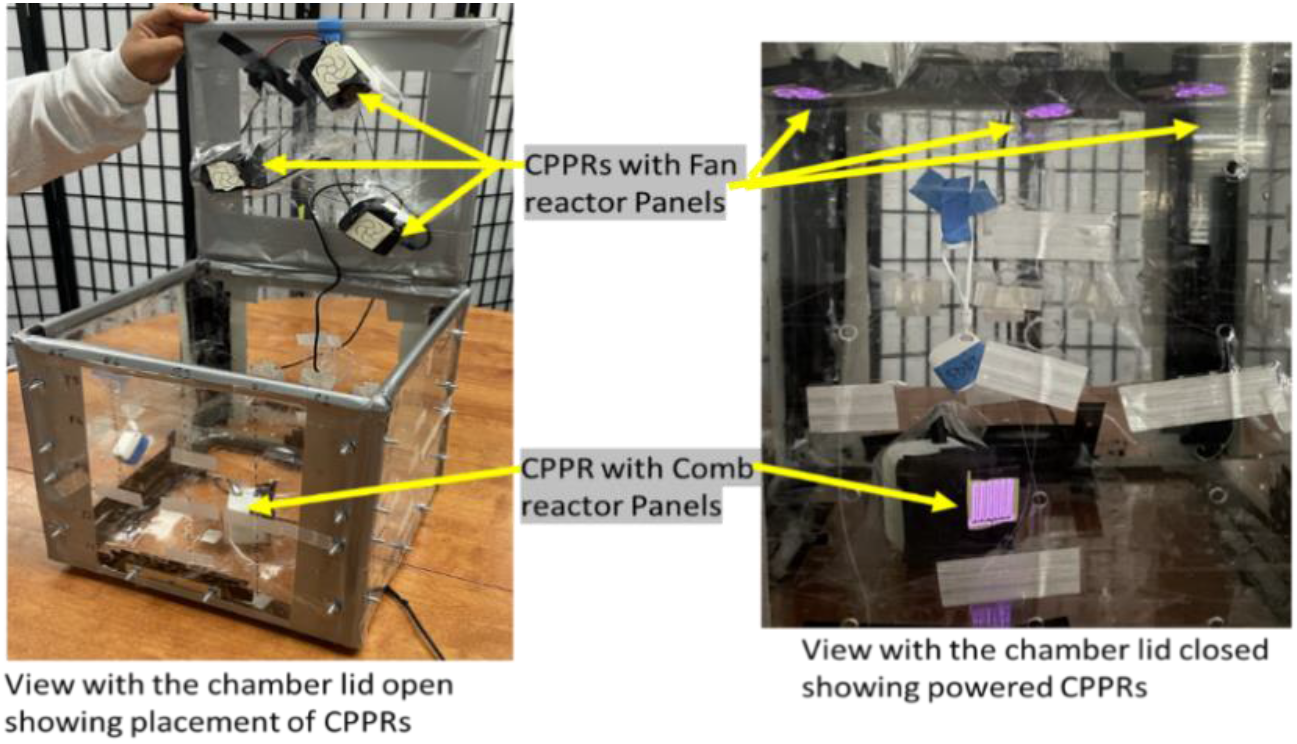
Integration of CPPRs and their placement inside the sterilization box.

### Evaluation of efficacy of the APS in sterilizing contaminated materials of interest

Experiments were performed to test sterilization efficacy of the developed APS prototype and included (a) 11 measurement points in the sterilization box, (b) two test organisms commonly used for sterilization tests (c) four materials commonly used in spacecraft applications (d) up to 6 CPPRs to determine minimum number of required CPPRs to obtain sterilization within 30 minutes (e) 7 exposure timepoints. Each of these components are described below, followed by the experimental procedure.

#### Measurement grid

11 measurement points were chosen in the internal volume of the prototype to simulate distributed decontamination of various points on the surface of an object placed inside the box. This included 9 points on Plane P2 (central plane), and center of planes P1 and P3 as shown in Figure 4. For suspending coupons inside the chamber in the measurement grid, sterile Teflon coated strings (0.1 mm diameter) were used to avoid ozone loss as Teflon does not react with ozone.

**Figure 4:**
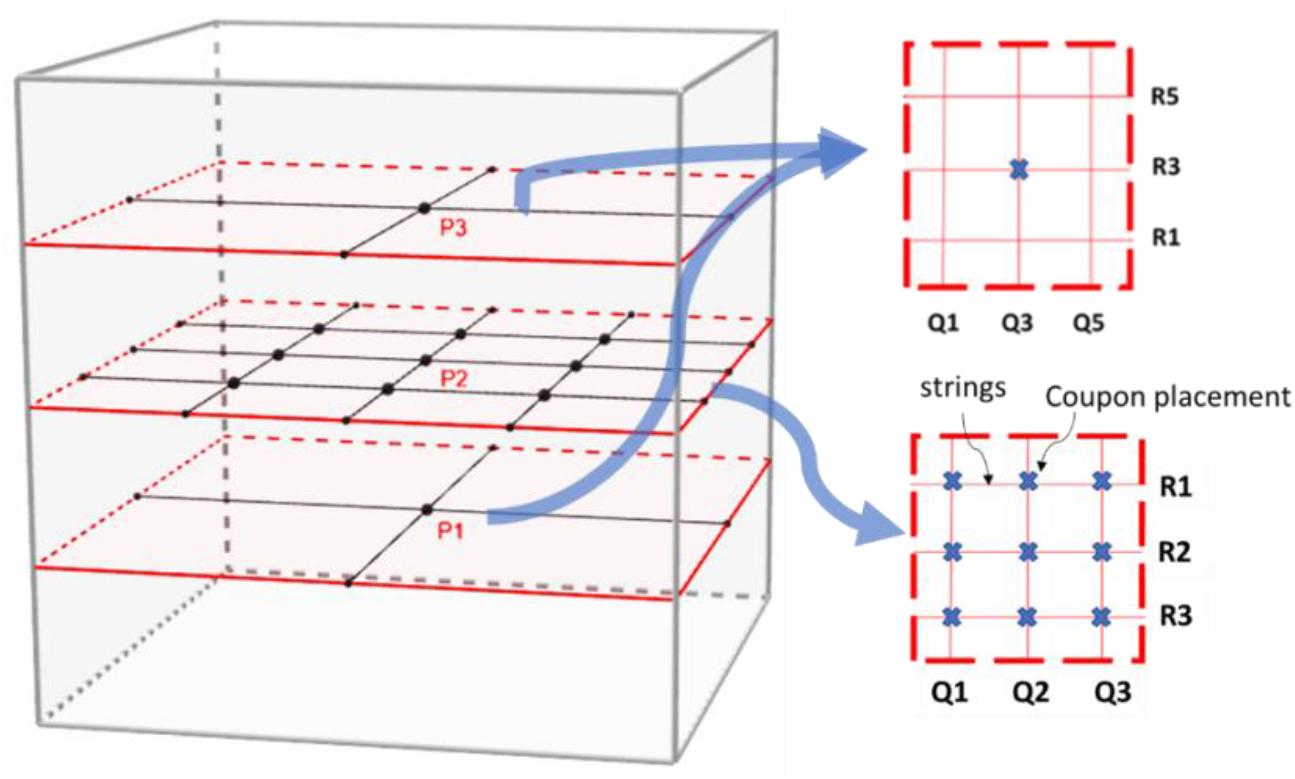
Measurement grid for decontamination measurement in three planes

#### Test organisms

*Escherichia coli* and *Bacillus subtilis* were the test organisms used in this study based on the proposed biological challenge list in Mars sample return planning document [24] and are described below.

*Escherichia coli* (ATCC 11775) NCTC 9001 *Escherichia coli* (Migula) Castellani and Chalmers, Serovar O1:K1:H7, Type strain. ATCC® 11775 (BSL 2 level): *Escherichia coli* (*E. coli*), a gram-negative bacterium, is a rod-shaped, facultative anaerobe with a replication ability under unfavorable conditions. This ability makes this species suitable for evaluation of decontamination technologies [26].

*Bacillus subtilis* (ATCC 6051) (Ehrenberg) Cohn, Type strain, Bacteriophage host (BSL 1 level): *Bacillus subtilis* (*B. subtilis*), is a spore forming gram positive aerobe generally found in soil and vegetation. Although non-pathogenic, it can contaminate food and be pathogenic for immuno-compromised people. This bacterial species was chosen as it is widely used in disinfection studies employing traditional disinfection methods [27]. Moreover, due to its spore forming ability, this species is of interest in evaluating decontamination technologies for planetary protection.

#### Test materials

Four test materials were chosen to be tested based on their important usage in space missions [6, 23]. These materials were cut to size 1 square inch to give coupons of four materials. The materials are presented in the table below with referenced example usage:

#### No. of CPPRs and exposure times

Exposure times refer to the period for which the inoculated coupons were placed inside the APS prototype. We iteratively tested the following combination of number of CPPRs and exposure times (CPPR ON + CPPR OFF times) to determine the number of CPPRs required to achieve complete killing of chosen bacteria species.

- 3 CPPRs: 10 mins (ON throughout).
- 4 CPPRs: 5 mins (ON throughout).
- 4 CPPRs: 15 minutes (10 mins ON + 5 mins OFF)
- 4 CPPRs: 20 minutes (15 mins ON + 5 mins OFF)
- 4 CPPRs: 25 minutes (20 mins ON + 5 mins OFF)
- 4 CPPRs: 30 minutes (15 mins ON + 5 mins OFF +5 mins ON + 5 mins OFF)
- 6 CPPRs - 30 minutes (25 mins ON + 5 mins OFF)

#### Preparation of Cultures

Stocks of *E.coli* and *B. subtilis* strains were stored at −80°C in LB (Luria-Bertani) broth and Nutrient broth, respectively, with 30% glycerol. Frozen stocks of *E.coli* and *B. subtilis* were grown overnight in LB broth at 37°C and Nutrient broth at 30°C, respectively. A Spectrophotometer was used for estimating the concentration of bacteria in the fresh LB and nutrient broth culture, followed by dilution, if necessary, to get approximately 5×10^7^ CFU (colony forming units)/ml. 10 μl of these broth cultures was used to inoculate coupons of the selected materials with 4 to 5 logs of *E.coli* and *B. subtilis*.

#### Pre-processing of coupons and APS box

The coupons were sterilized by autoclaving them at 121°C in a dry autoclave cycle. In the beginning of every experiment, the box and all the components inside it were wiped with 70% isopropyl alcohol to avoid external contamination. Further, it was ensured that ozone concentrations inside the box prior to starting an experiment matched room level ozone concentrations by giving at-least half an hour of free air flow inside the box with the lid open. This is done by placing the chamber in a ducted BYPASS fume hood – Phoenix Controls Corporation – 100 lfm. All experiments were conducted in this fume hood for safety. Note that the box was sealed during experiments to ensure no effect of the fume hood during an experiment.

#### Post-processing of coupons and APS box

Post experiments, the coupons were mixed thoroughly in 15 ml PBS solution using a Fisher Scientific ® Mini Vortexer lab mixer and 100ul of this mixture (for exposed coupons) or its dilution (for unexposed coupons - Control) was plated on agar plates (LB agar for *E. coli* and Nutrient Agar for *B. subtilis*) followed by incubation at 37°C (*E. coli*) and 30°C (*B. subtilis*) for 24 hours. Plate counts were obtained to quantify the bacterial colonies present in the coupons. All post processing was conducted in a Biological Safety Cabinet (BSC) Class II, Type A2 to avoid external contamination and maintain safety protocols.

#### Control Experiments

Control experiments were performed to ensure correct quantification of inactivation of bacteria due to the APS and rule out inactivation due to environmental factors related to the experimental setup or procedure. This involved establishing consistent bacterial concentrations for the sterilized coupons inoculated with a fixed volume of the bacterial culture. For these experiments, inoculated coupons were left inside the APS for periods matching exposure times without powering up the CPPRS and post processed. At least three repeats were performed.

#### Exposure Experiments

Each exposure experiment involved 14 coupons of one material inoculated with 10 μl of bacteria culture containing approximately 10^7^ CFUs/ml of one type of bacteria. Inoculation volume of 10 μl was chosen to give 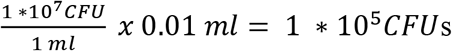 i.e. approximately 10^5^ CFUs/coupon. 11 coupons were placed in the APS at the 11 measurement points shown in Figure 4 and exposed for selected times. The remaining 3 coupons were placed outside the chamber for the same times as controls. After the exposure periods, all 14 coupons were post processed to obtain CFUs/ coupon. The following equation was used to calculate CFUs/coupon:

CFUs/coupon = CFUs/ml * V1 = *Dx* * 10^x^ * *V*1; V1 = volume of PBS used to mix coupons in post processing in mls, and Dx = CFUs counted in x^th^ dilution plate.

The reduction in microbial colonies obtained per coupon at each measurement point was determined from the difference in CFUs/coupon of the exposed and control (unexposed) coupons.

### Ozone measurements

The 2B Technologies Model 106-6 Ozone Monitor™, which works based on UV light absorption at 254 nm, was used for the ozone measurements in the APS prototype chamber [28]. The accuracy of the monitor is 0.01 ppm or 2% of the reading. Ozone measurements were performed at the center of the sterilization box for the exposure times determined in the sterilization efficacy tests with corresponding CPPR ON and OFF times.

### Ozone penetration tests

An enclosure was built to test CPPR generated ozone permeability through fabric layers without the help of an external agent. As shown in Figure 5, the enclosure has the following components: 1) CPPR support tabs: to hold the CPPR; 2,3) top and bottom measurement holes: positioned above and below the fabric layer for access to data collecting probe; 4) air exit hole: this is connected to left open, 5) Fabric sample holder: to hold the fabric samples or cells. A CPPR with the fan reactor panel was used for these studies. The CPPR was held using the CPPR support tabs such that the fan reactor panel positioned inside the enclosure is facing down. This was done to employ the flow produced by the fan reactor for pushing the ozone through the fabric layer samples. The fabric sample holder had a rectangular cutout with Velcro attached to attach to Velcro on the boundary of the fabric samples. Ozone concentrations were measured above and below the fabric layers for 5 minutes after 10 seconds of powering up the CPPRs to determine ozone permeability. Measurements were averaged over the last minute of the measurement period when ozone concentrations stabilized. Top and bottom measurements were performed separately to avoid effect of measurement on data collection. The top and bottom of the enclosure was blocked to mitigate generated ozone from escaping. Before every experiment it was ensured that ozone levels inside the enclosure was equal to ambient ozone levels. Three repeats were performed for each fabric sample type.

**Figure 5.**
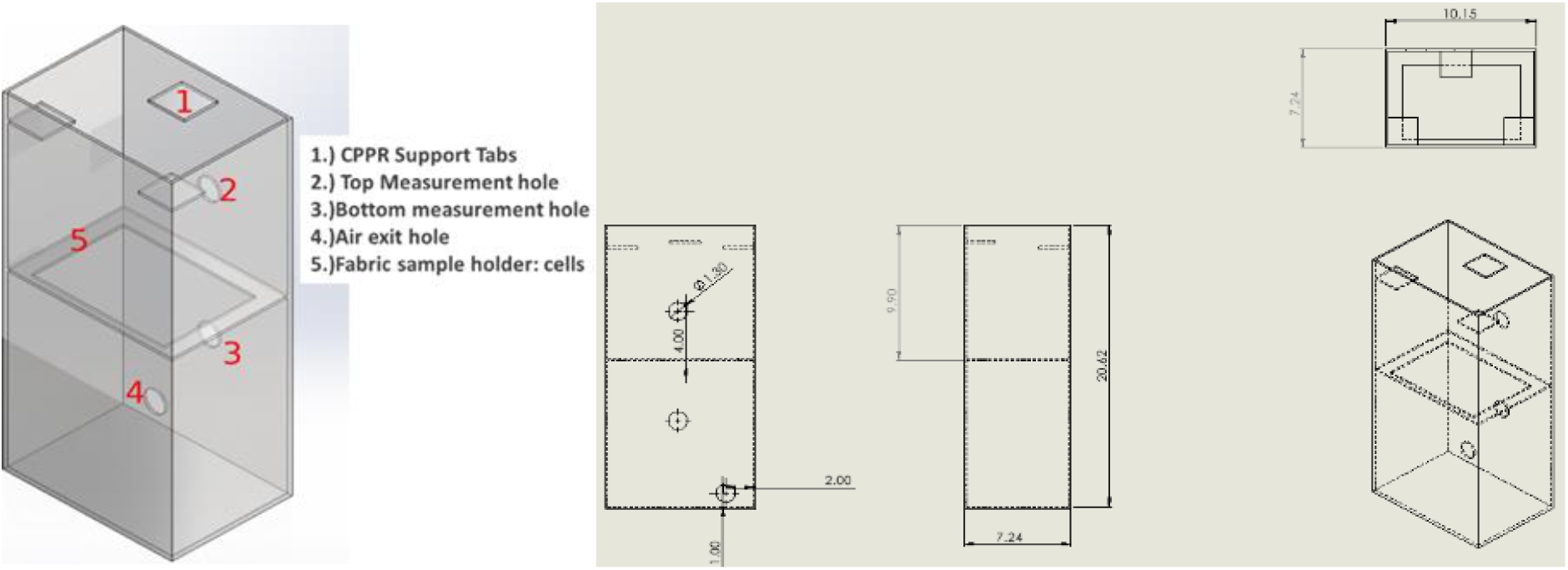
Schematic and dimensions of the enclosure built for ozone penetration tests.

Three fabric types were tested based on their use in planetary protection studies [23]: Kevlar, Orthofabric and a combined layer of Kevlar and Orthofabric. Ozone data was collected before and after the fabric layers. The samples were sized 4 cm x 5 cm with thickness of 1 mm, 2 mm and 3 mm for Orthofabric, Kevlar and combined samples, respectively. The Kevlar fabric had a thickness of 2mm meanwhile the Orthofabric had a thickness of 1mm. The samples were held in “cells” made of Velcro which hold the fabric and keep it in a uniform shape as shown in Figure 6.

**Figure 6:**
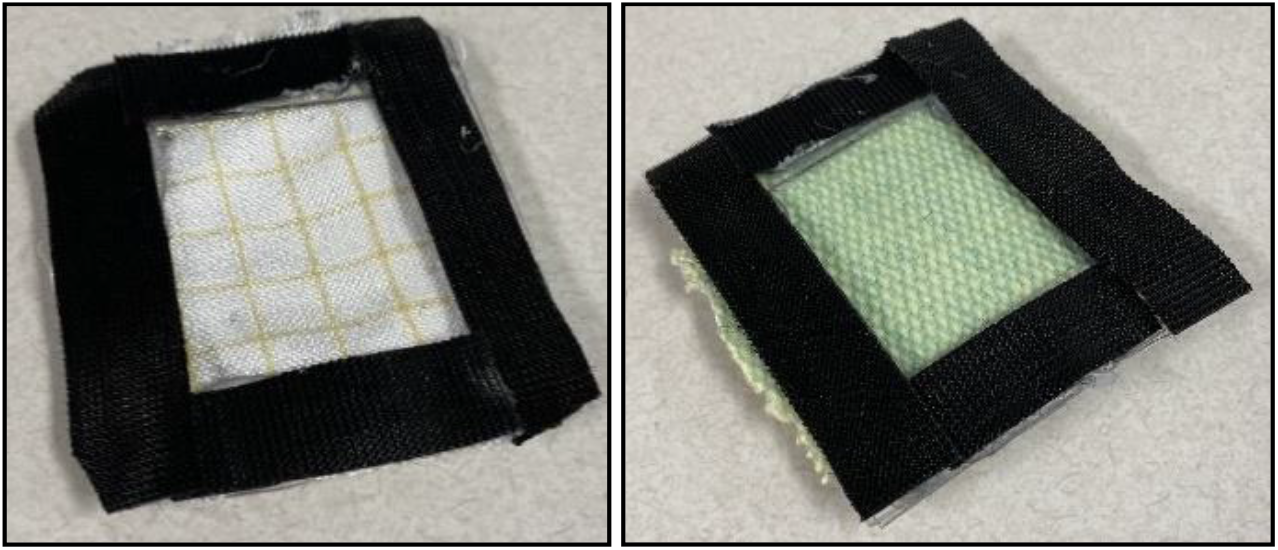
Orthofabric(left) and Kevlar(right) Fabric Cells.

Ozone was measured using the 106-M Ozone Monitor by 2B Technologies as mentioned before. Temperature was measured using a Testo 405i Smart Probe Hotwire anemometer with accuracy ±0.5 °C of measured temperature.

### Preliminary material compatibility tests

The four selected materials: Aluminum, Polycarbonate, Kevlar and Orthofabric, were exposed in the APS for exposure conditions that resulted in complete killing of *E. coli* and *B. subtilis*. These exposure conditions correspond to equivalent ozone CT (concentration x time) values required for sterilization (4 to 5 log reduction) of the two test species. All four materials were tested for visible surface degradation and change in material composition using standard SEM analysis. The SEM Hitachi S 3000, at the Nanoscale Research Facility (NRF) at the University of Florida was used to perform these tests [29].

## Results and Discussion

### Evaluation of efficacy of the APS in sterilizing contaminated materials of interest

#### Control experiments

The results obtained for establishing consistent bacterial concentrations for the sterilized coupons of different material inoculated with 4 to 6 logs of *E. coli* and *B. subtilis* CFUs are shown in Figures 7 and 8 with error bars based on standard deviation from 3 repeats. The data shows that inoculating same sized coupons of same material with same amount of culture resulted in consistent bacterial count per coupon with a maximum variation of 0.3 log10 (CFU/coupon). This data also validated the post processing method to recover microbial population from inoculated coupons.

**Figure 7:**
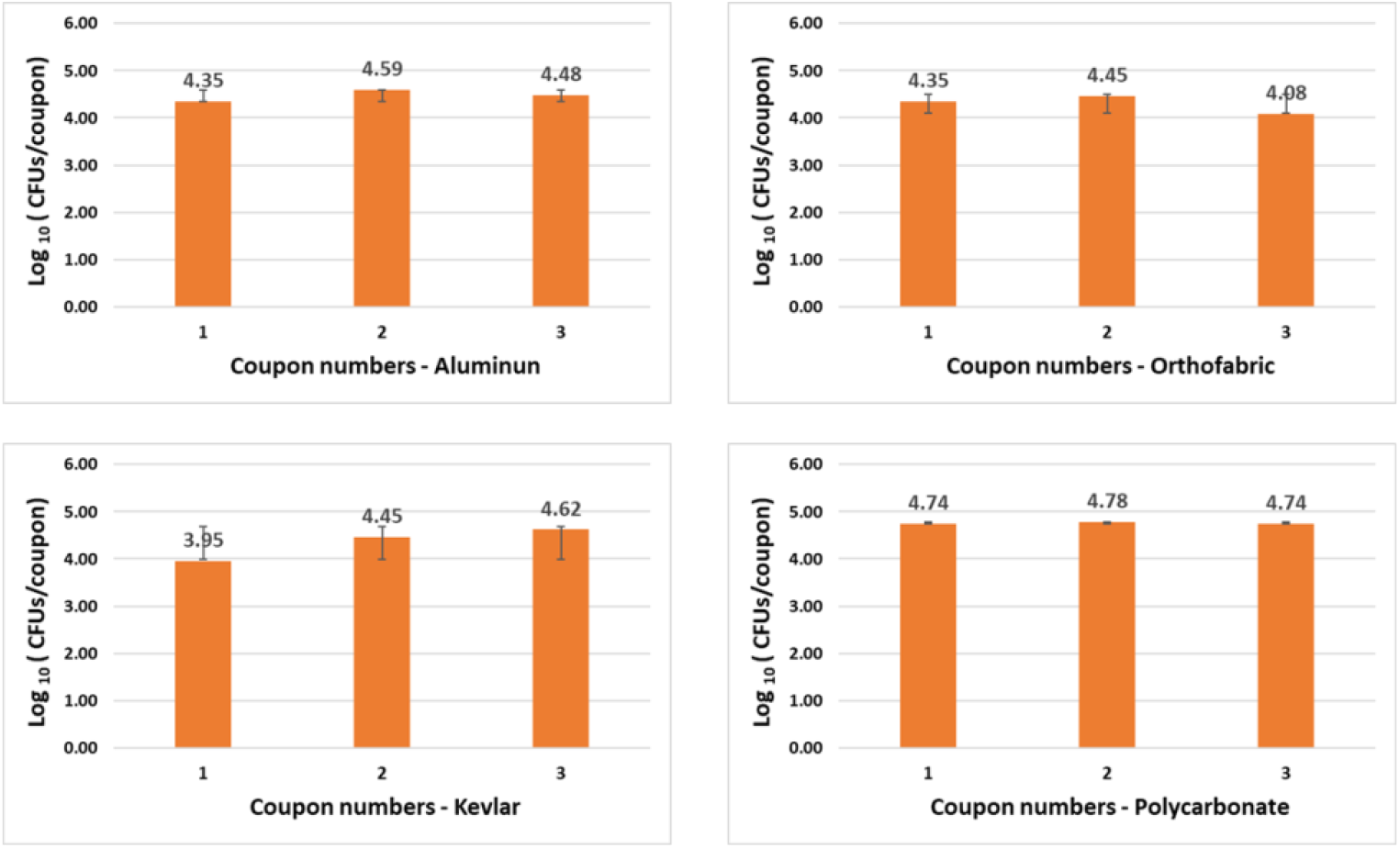
Data establishing consistent controls with E. coli inoculation, i.e., inoculating coupons of same material with same volume of E. coli culture resulted in consistent bacterial count per coupon with a maximum variation of 0.3 log10 (CFU/coupon).

**Figure 8:**
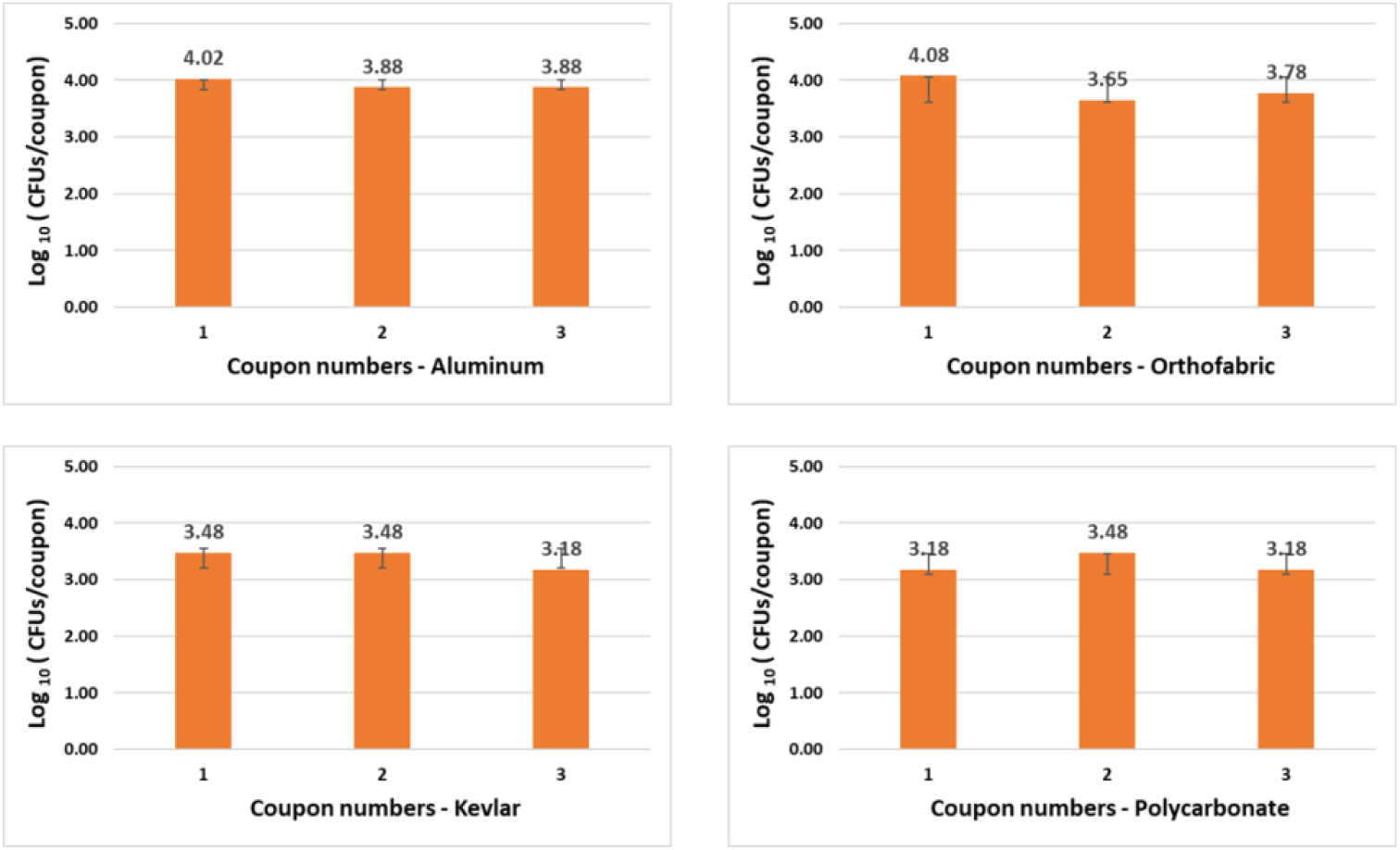
Data establishing consistent controls with B. subtilis inoculation, i.e., inoculating coupons of same material with same volume of B. subtilis culture resulted in consistent bacterial count per coupon with a maximum variation of 0.3 log10 (CFU/coupon).

#### Exposure experiments

A total of 37 iterative exposure experiments were performed with two test organisms, 7 exposure time points and 4 selected materials as mentioned in the Materials and Methods section. The goal of the iterative experiments was to find the optimum combination of no. of CPPRs and exposure time required for complete killing of each test organisms for all selected material within 30 minutes. Results showed that with a control count of 4 to 5 logs of CFU/coupon, complete killing for *E. coli* was achieved with 4 CPPRs in 20 minutes. The same was achieved for *B. subtilis* with 6 CPPRs in 30 minutes. Figures 9 to 12 below show the results of the overall sterilization (average reduction over 11 points of measurement) and spatial distribution of log reductions achieved for each material type contaminated with *E. coli* and *B. subtilis* and subjected to APS exposures corresponding to (a) 20 minutes exposure time with 4 CPPRs (15 mins ON + 5 mins OFF) and (b) 30 minutes exposure time with 6 CPPRs (25 mins ON + 5 mins OFF). All experimental data from the iterative experiments can be found in supplementary materials.

**Figure 9:**
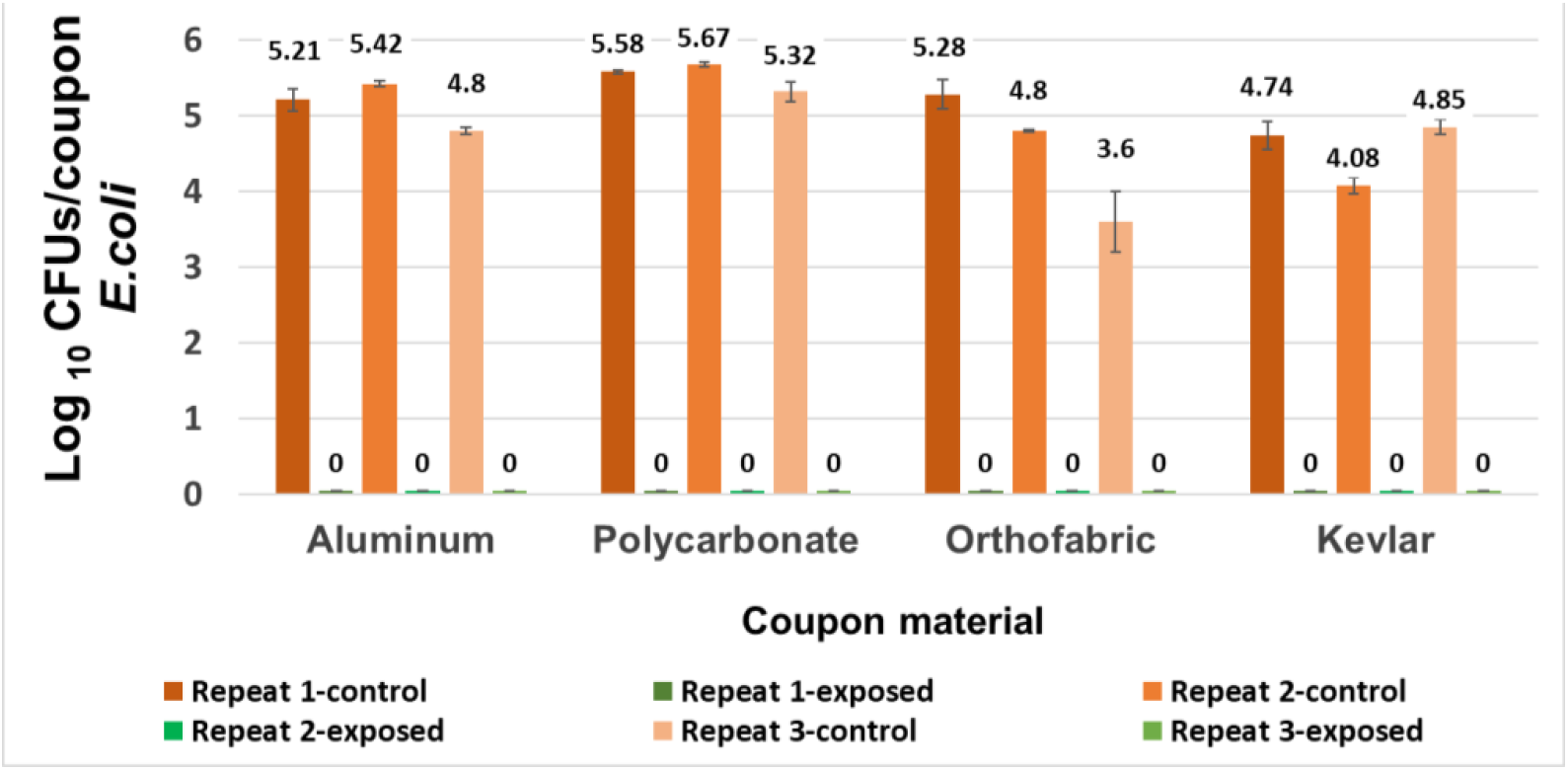
Data showing logs of E. coli CFUs/coupon of exposed coupons (averaged over 11 coupons placed inside the APS) and control coupons (averaged over 3 coupons placed outside the APS) for four coupon materials, each repeated three times. Complete killing (4 to 5 log reduction) was achieved in each case using 4 CPPRs and exposures of 20 minutes (15 mins CPPR on + 5 mins CPPR off).

**Figure 10:**
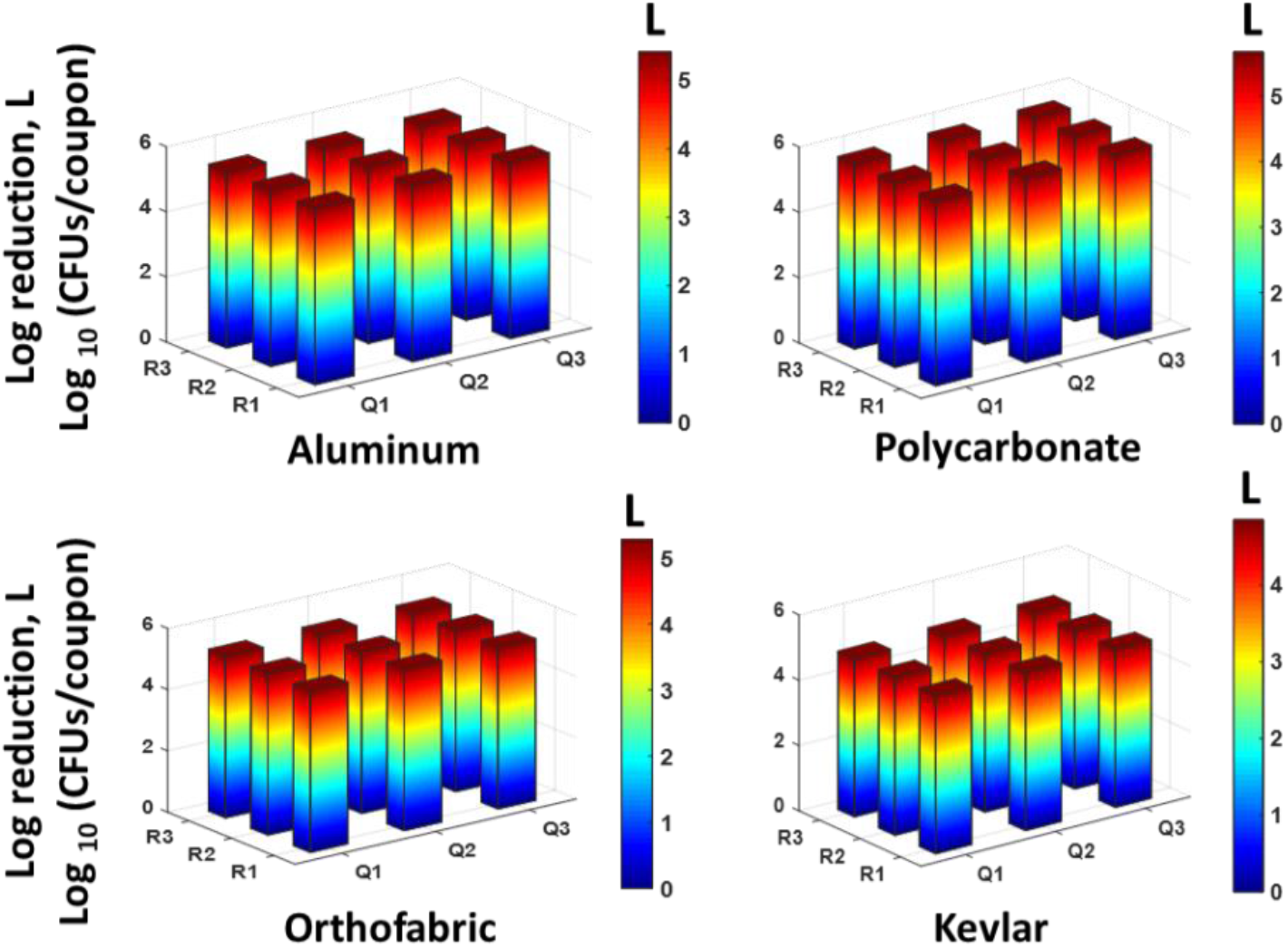
Data showing uniform spatial distribution of reduction in logs of E. coli CFUs/coupon of 9 exposed coupons placed at the central plane of the APS, for each material type. Complete killing (4 to 5 log reduction) was achieved at all points using 4 CPPRs and exposures of 20 minutes (15 mins CPPR on + 5 mins CPPR off)

**Figure 11:**
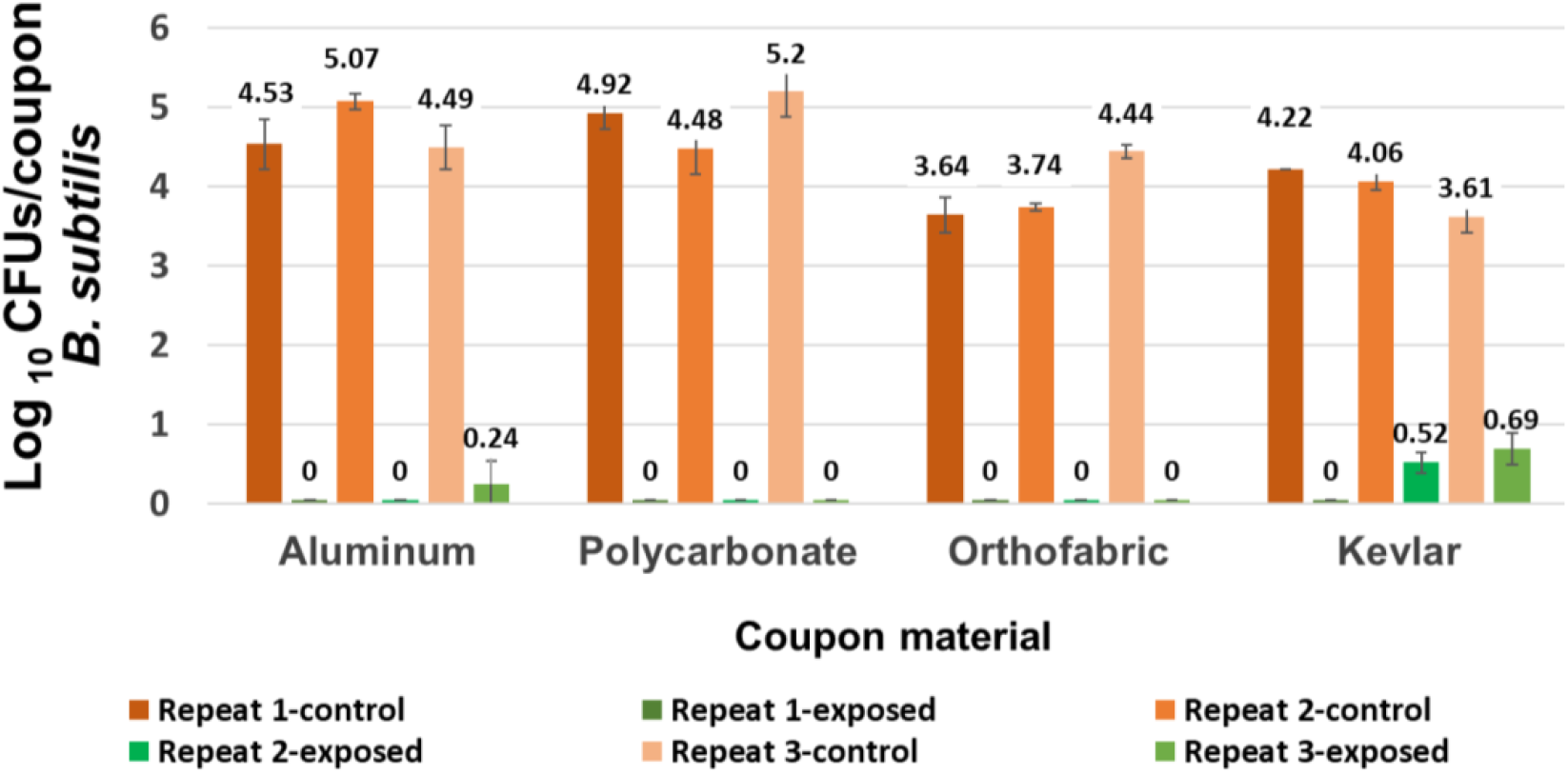
Data showing logs of B. subtilis CFUs/coupon of exposed coupons (averaged over 11 coupons placed inside the APS) and control coupons (averaged over 3 coupons placed outside the APS) for four coupon materials, each repeated three times. Complete killing (4 to 5 log reduction) was achieved in most cases using 6 CPPRs and exposures of 30 minutes (25 mins CPPR on + 5 mins CPPR off)

**Figure 12:**
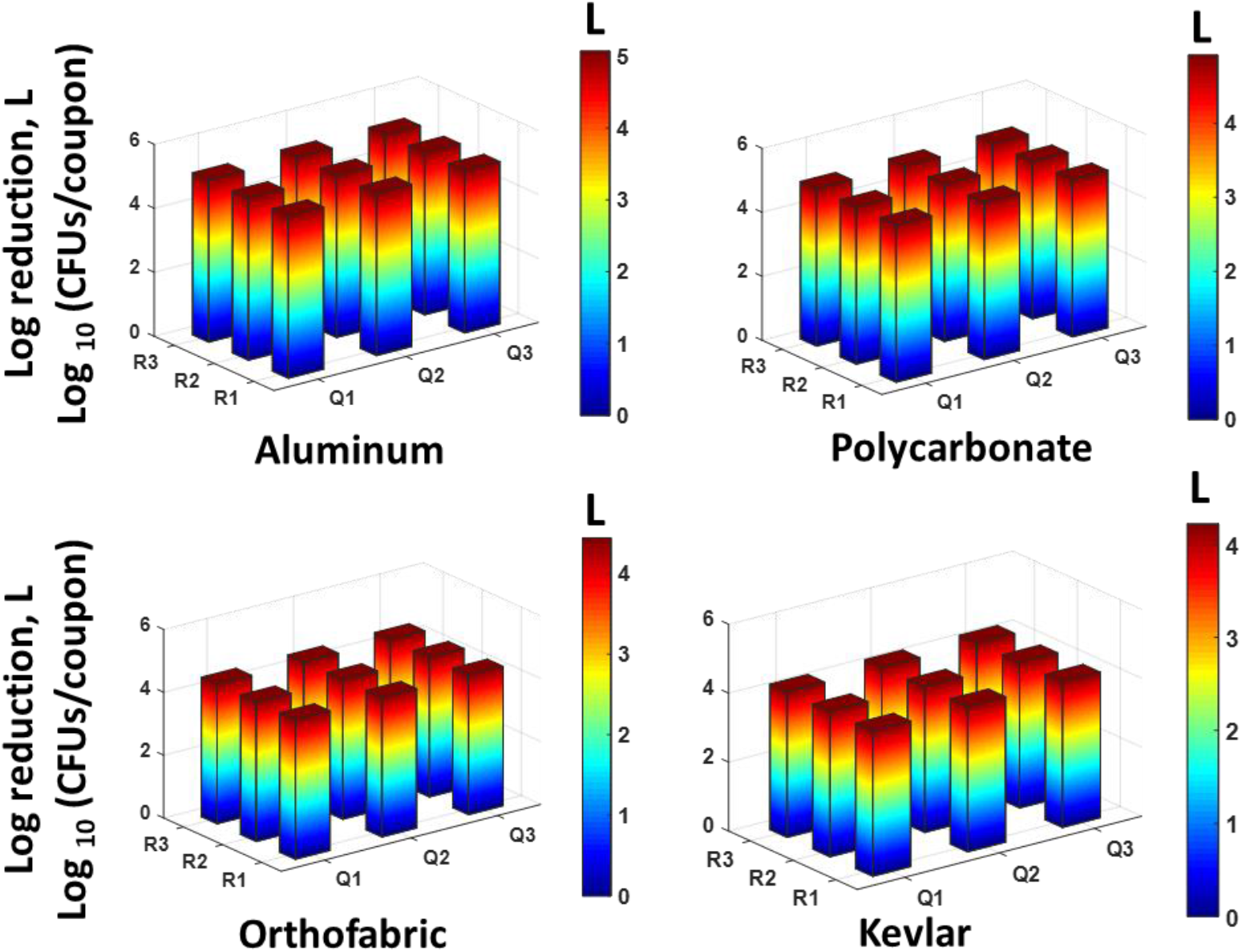
Data showing uniform spatial distribution of reduction in logs of B. subtilis CFUs/coupon of 9 exposed coupons placed at the central plane of the APS, for each material type. 4 to 5 log reduction was achieved at all points using 6 CPPRs and exposures of 30 minutes (25 mins CPPR on + 5 mins CPPR off).

The results obtained for *E. coli* contaminated surfaces (Figures 9 and 10) show that the APS achieved complete killing or 4 to 5 log reductions of *E. coli* on Aluminum, Polycarbonate, Orthofabric and Kevlar, at 11 points inside the chamber, with a minimum exposure time of 20 minutes with 4 CPPRs active for 15 minutes. Similarly, figures 11 and 12 show that the APS achieved complete killing of *B. subtilis* on Aluminum, Polycarbonate, Orthofabric and Kevlar, at 11 points inside the chamber, with a minimum exposure time of 30 minutes with 6 CPPRs active for 25 minutes. Compared to *E. Coli*, additional CPPRs and exposure times required for complete killing of *B. subtilis* can be explained by *B. subtilis* being a Gram – positive bacteria: it lacks an outer membrane as Gram-negative but are surrounded by layers of peptidoglycan many times thicker than is found in the Gram-negatives, with protective cell wall outside the cell membrane - it has a cytoplasmic membrane and a thick cell wall. Further, as a spore-forming bacteria, the selected *B. subtilis* strain can form a tough, protective endospore for resistance, allowing it to tolerate extreme environmental conditions.

The sterilization efficacy results show the potential of APS in uniformly sterilizing an object made of Aluminum, Orthofabric, Polycarbonate or Kevlar, contaminated with 4 to 5 logs of pathogens like *E. coli* and *B. subtilis*, per square inch of surface, within 30 minutes. Further, successful sterilization achieved for the four materials establishes that the APS is capable of sterilizing both solid and porous surfaces without the need of an external mixing or distributing agent.

### Ozone data and CT (concentration x time) requirements

Based on the number of CPPRs and corresponding exposure times that resulted in complete killing of 4 to 5 logs of *E. coli* and *B. subtilis* concentrations on 11 coupons inside the APS chamber, ozone data was collected in the APS at the center point of the central plane (P2) for the following operating conditions (a) 4 CPPRs and 20 minutes (15 min CPPRs on, 5 mins CPPRs off), (b) 6 CPPRs and 30 minutes (25 min CPPRs on, 5 mins CPPRs off). These measurements were performed separate from the decontamination experiments to (a) avoid incorrect measurements due to loss of concentrations measured because of usage of ozone for decontamination and (b) avoid contamination of the ozone monitor. The results are shown in Figure 13 with the orange line representing the time when the CPPRs were turned off. Three repeats were performed for each exposure time to gain statistical confidence and standard deviation observed in those repeats were used to represent error bars in the graph.

**Figure 13:**
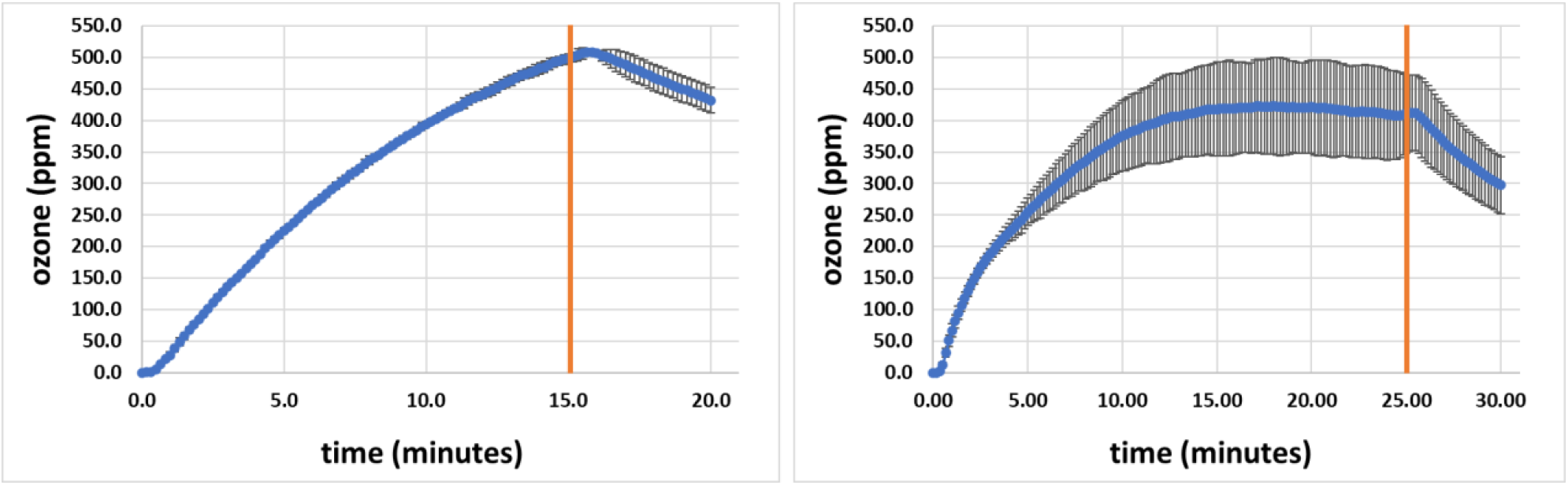
APS ozone data for (a) 4 CPPRs and 20 minutes exposure (15 min CPPRs on, 5 mins CPPRs off) and (b) 6 CPPRs and 30 minutes exposure (25 min CPPRs on, 5 mins CPPRs off)

Figure 13(a) shows that the ozone concentrations reach a peak of around 500 ppm in 15 minutes at the center of the APS and gradually decrease thereafter when 4 CPPRS were powered for 20 minutes (15 mins ON + 5 mins OFF). Interestingly, when 6 CPPRS were powered 30 mins (25 mins ON + 5 mins OFF) the ozone concentrations at the center of the APS converge to around 420 ppm in 15 minutes and stabilize at that concentration till the CPPRs are turned off. Although this seems counter-intuitive, possible explanations for this difference can be that (a) with 6 CPPRs distributing the generated ozone, more mixing of ozone occurs inside the APS which can help to better decontaminate a larger volume but can also affect ozone measurements or result in faster decomposition of ozone, and (b) the peak concentration at the center of the APS might shift to another point with more mixing and vortical structures formed due to interaction of flow structures generated by the 6 CPPRs. Furthermore, increased deviations observed in the concentrations with 6 CPPRs powered up in the APS can be attributed to increased mixing and unstable flow structures. It is to be noted here that measurements of spatial ozone distribution inside the APS was out of scope for this study due to limitation of time and more focus on spatial distribution of decontamination inside the APS. A detailed study of spatial and temporal ozone distribution by SDBD reactors can be found in a recent study published in this area [15].

Ozone CT requirements are calculated using the following formula to account for difference in CPPR on and off times during an exposure:

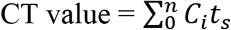

Where C_i_ refers to i^th^ sample reading given by the ozone monitor, t_s_ refers to sampling time of the ozone monitor and n is the total number of samples collected during a specific exposure time. The calculated CT values along with maximum ozone concentrations achieved for each exposure time is given below in Table 2.

**Table 1:**
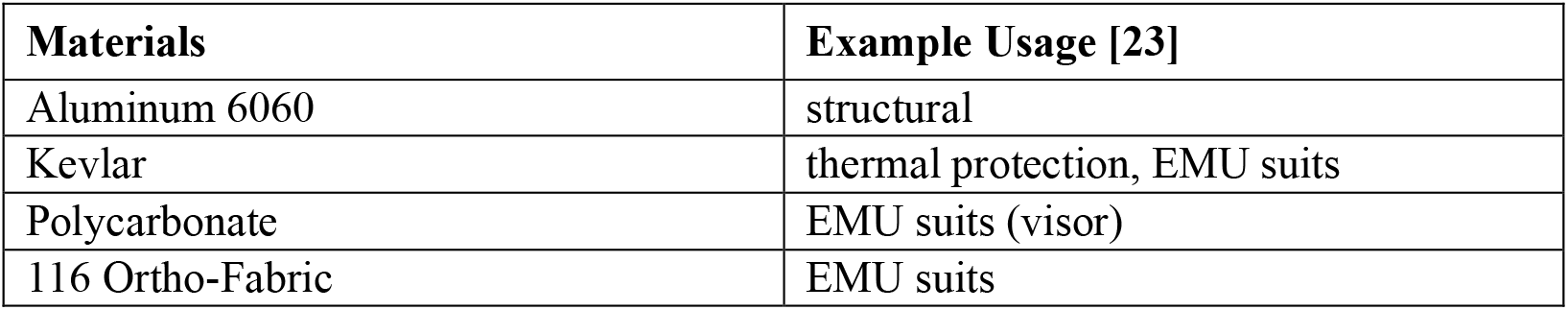
Materials selected to be tested:

**Table 2:** Ozone CT (concentration x time) values in APS for the number of CPPRs and exposure times.

| Exposure time (minutes) | No. of CPPRs | Ozone CT values (ppm-min) |
| --- | --- | --- |
| 20 (15 min CPPR on + 5 min CPPR off)<br>[required for 4-5 log reduction of *E. coli* on 4 selected materials] | 4 | 6789.02 |
| 30 (25 min CPPR on + 5 min CPPR off)<br>[required for 4-5 log reduction of *B.subtilis* on 4 selected materials] | 6 | 10323 |

### Power consumption

Based on the sterilization efficacy tests, 6 CPPRs used in the APS can achieve complete killing of 4 to 5 logs of pathogenic bacteria on various surfaces within 30 minutes with 6 CPPRs active for 25 minutes. With the power requirement of a single CPPR being 2.2+0.37 Watts [21], the power requirement of 6 CPPRS in the APS was calculated by multiplying by a factor of 6 giving total power consumption of 13.2 + 2.22 W.

### Ozone penetration

Averaged ozone data over 5 repeats of measurements above (upstream) and below (downstream) fabric layers is given in Figure 14. The percentage reduction in ozone concentration from above the fabric layer (the side where the CPPR was placed) to below the fabric sample was 7.59% for Kevlar, 16.17% for Orthofabric and 13.90% for a combined layer of Kevlar and Orthofabric. Average temperatures in the regions above and below the fabric ranged between 25.2 to 26.8°C which were close to measured room temperature of 24.4°C. These results indicate that the CPPR generated ozone can penetrate through the fabric layers in an enclosure without the aid of an external agent. Thus, the data shows that SDBD flow actuation can be used to facilitate ozone penetration and that an external pressure driven mechanism is not necessary for CPPR generated ozone penetration through fabric layers. This data combined with decontamination data for Kevlar and Orthofabric indicate that the APS will not require an additional component for ozone penetration through fabrics like Kevlar and Orthofabric.

**Figure 14.**
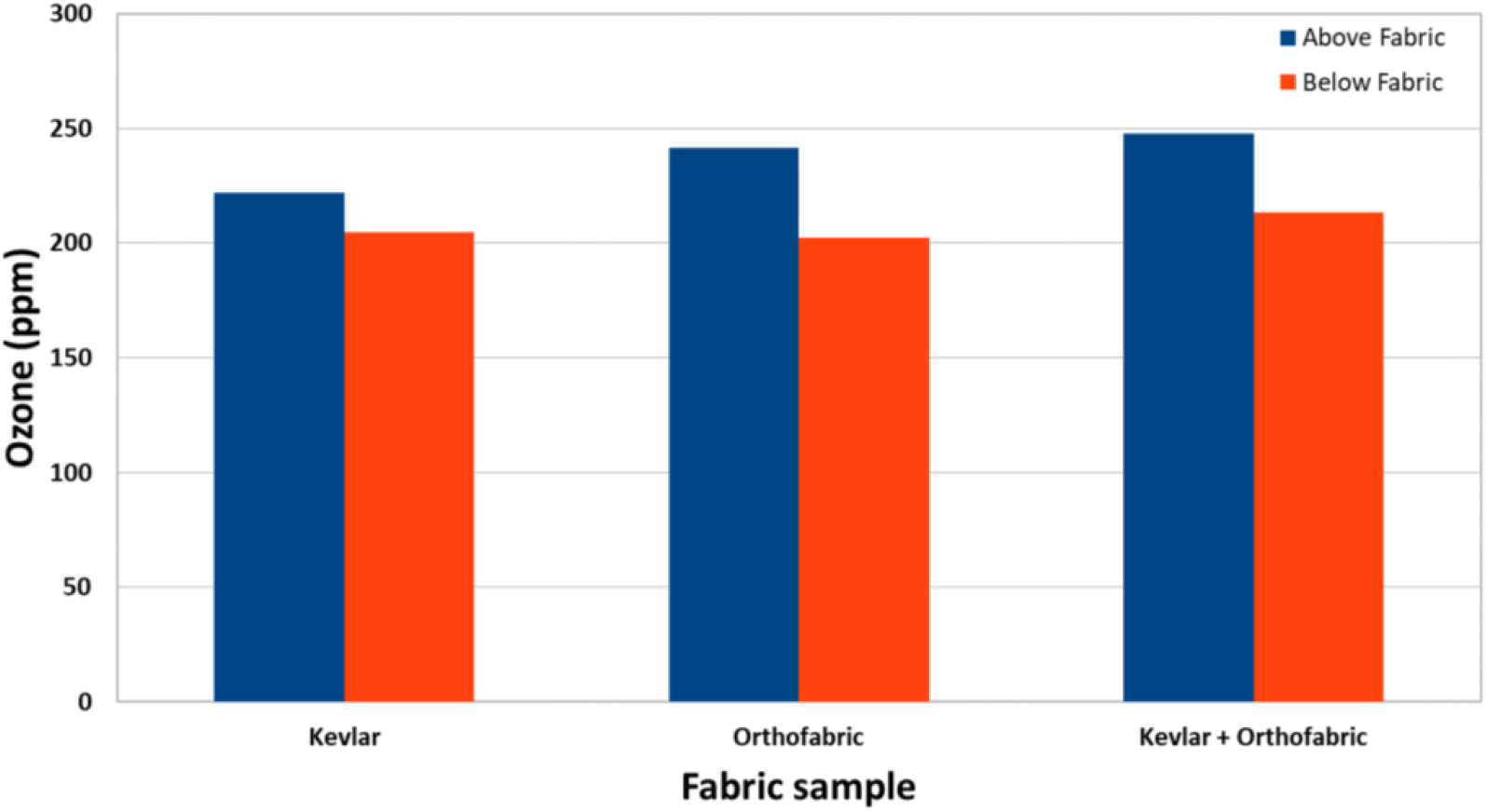
Ozone concentrations before and after fabric layers showing CPPR generated ozone penetration through fabric layer(s).

### Material Compatibility

One coupon of all four selected materials: Aluminum, Polycarbonate, Kevlar and Orthofabric, was exposed in the APS sterilization chamber for the two exposure conditions that resulted in complete killing of *E. coli* and *B. subtilis*, respectively: (a) APS Exposure 1: 20 minutes with 4 CPPRs (15 min on + 5 min off) i.e. exposed to ozone CT value of 6789.02 ppm-min and (b) APS Exposure 2: 30 minutes with 6 active CPPRs (25 min on + 5 min off) i.e. exposed to ozone CT value of 10323 ppm-min. All four materials were tested for visible surface degradation and change in material composition by comparing with a control coupon not exposed to the APS, using standard SEM analysis: results shown in Figures 15 to 18. Data for control (unexposed) material samples was compared with samples undergoing APS Exposure 1 and APS exposure 2. Visual analysis of the SEM images, at 50 to 200 microns scale, did not show material degradation. Comparison of material composition showed increase in oxygen content (by weight) by 2.8% for Aluminum, 0.1% for Polycarbonate, 0.9% for Orthofabric, and 3.1% for Kevlar for APS Exposure 1. Corresponding increase for APS Exposure 2 was observed to be 5.9% for Aluminum, 1.4% for Polycarbonate, 1.2% for Orthofabric, and 5.8% for Kevlar. These preliminary results suggest that APS exposures required for sterilization will not lead to considerable material degradation for Aluminum, Polycarbonate, Orthofabric and Kevlar material. Further SEM analysis with detailed compositional study will be performed in future studies of the APS.

**Figure 15.**
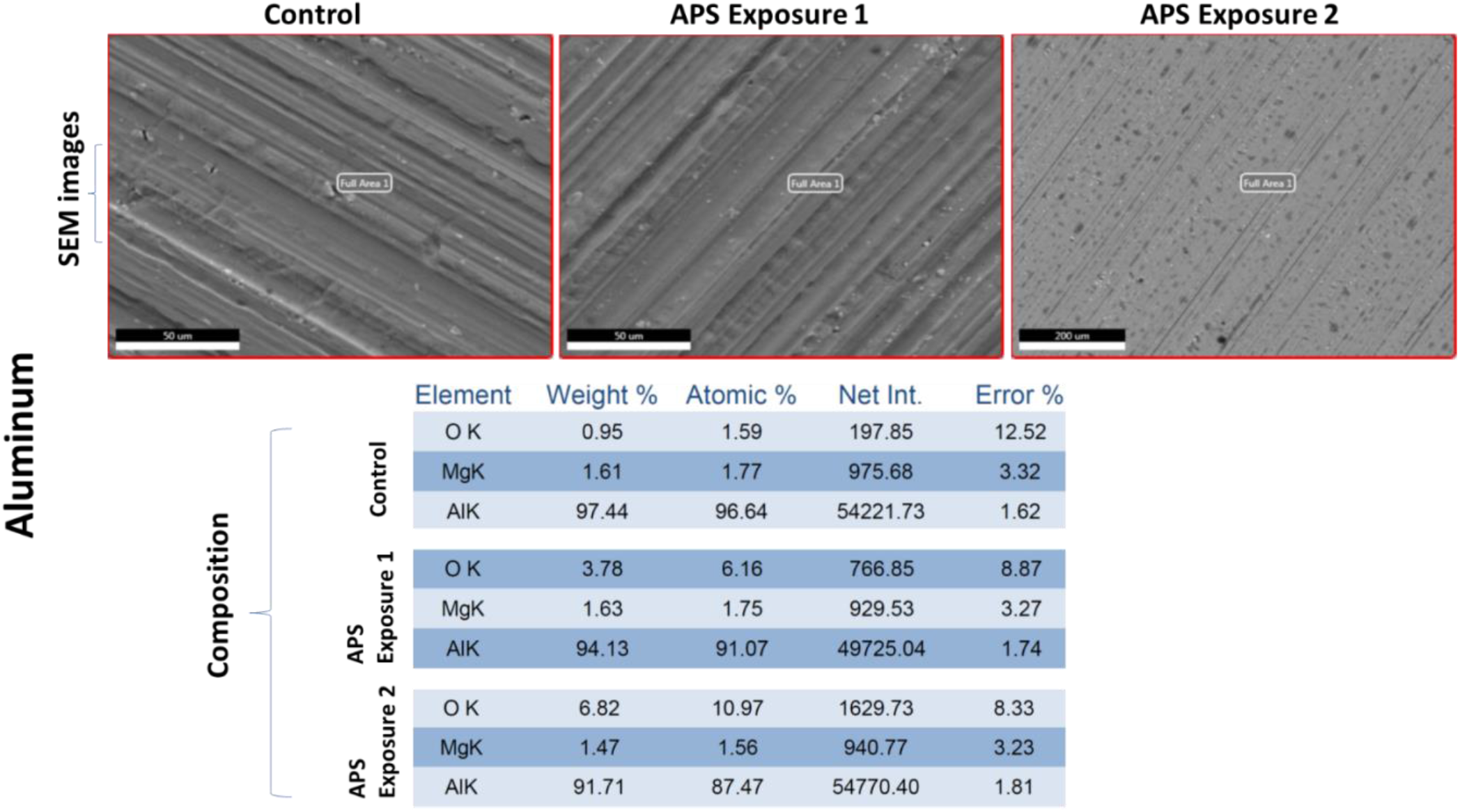
Material compatibility data: SEM analysis of Aluminum with APS exposure 1: exposed to ozone CT value of 6789.02 ppm-min, and APS exposure 2: exposed to ozone CT value of 10323 ppm-min.

**Figure 16.**
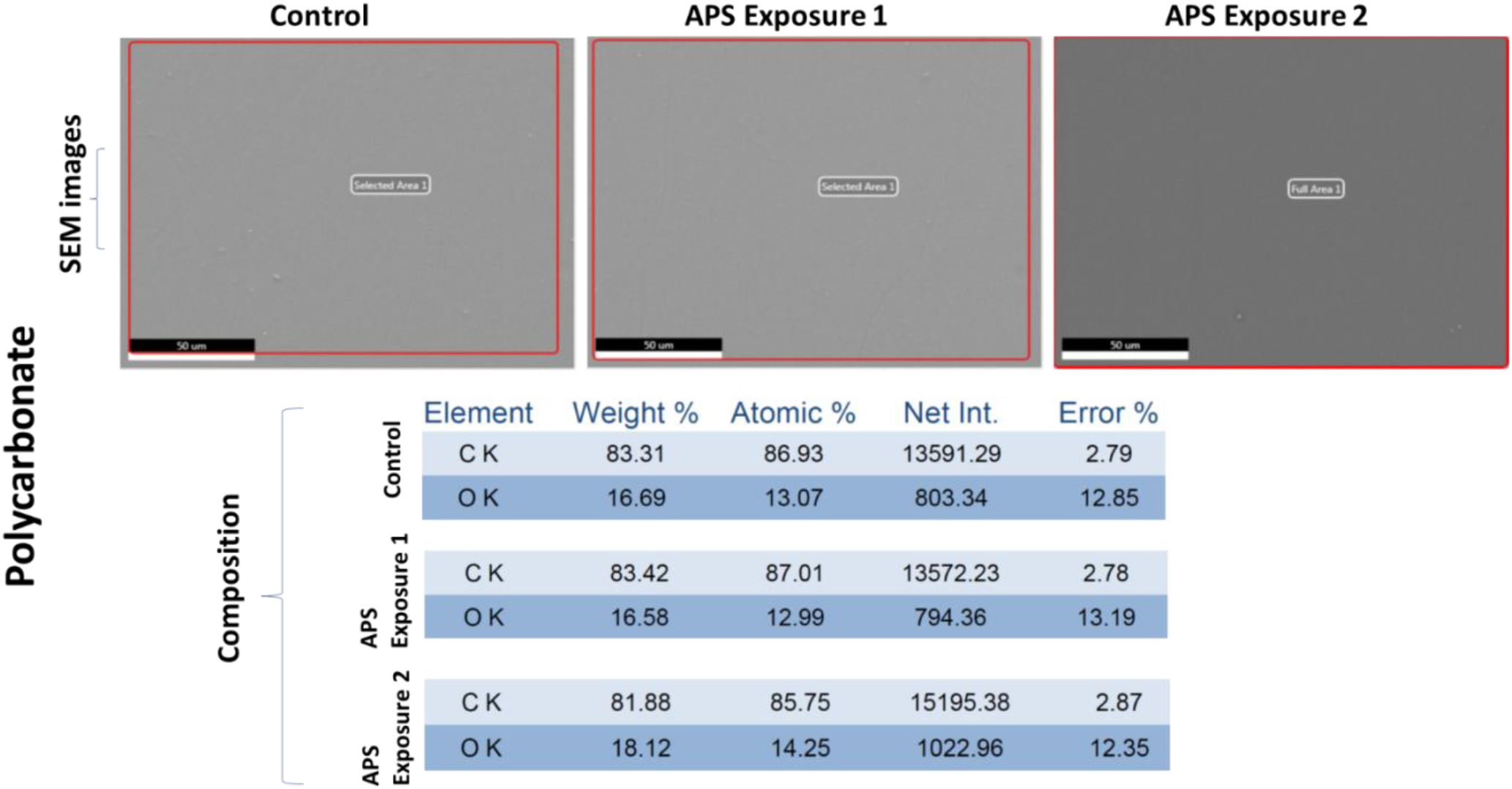
Material compatibility data: SEM analysis of Polycarbonate with APS exposure 1: exposed to ozone CT value of 6789.02 ppm-min, and APS exposure 2: exposed to ozone CT value of 10323 ppm-min.

**Figure 17.**
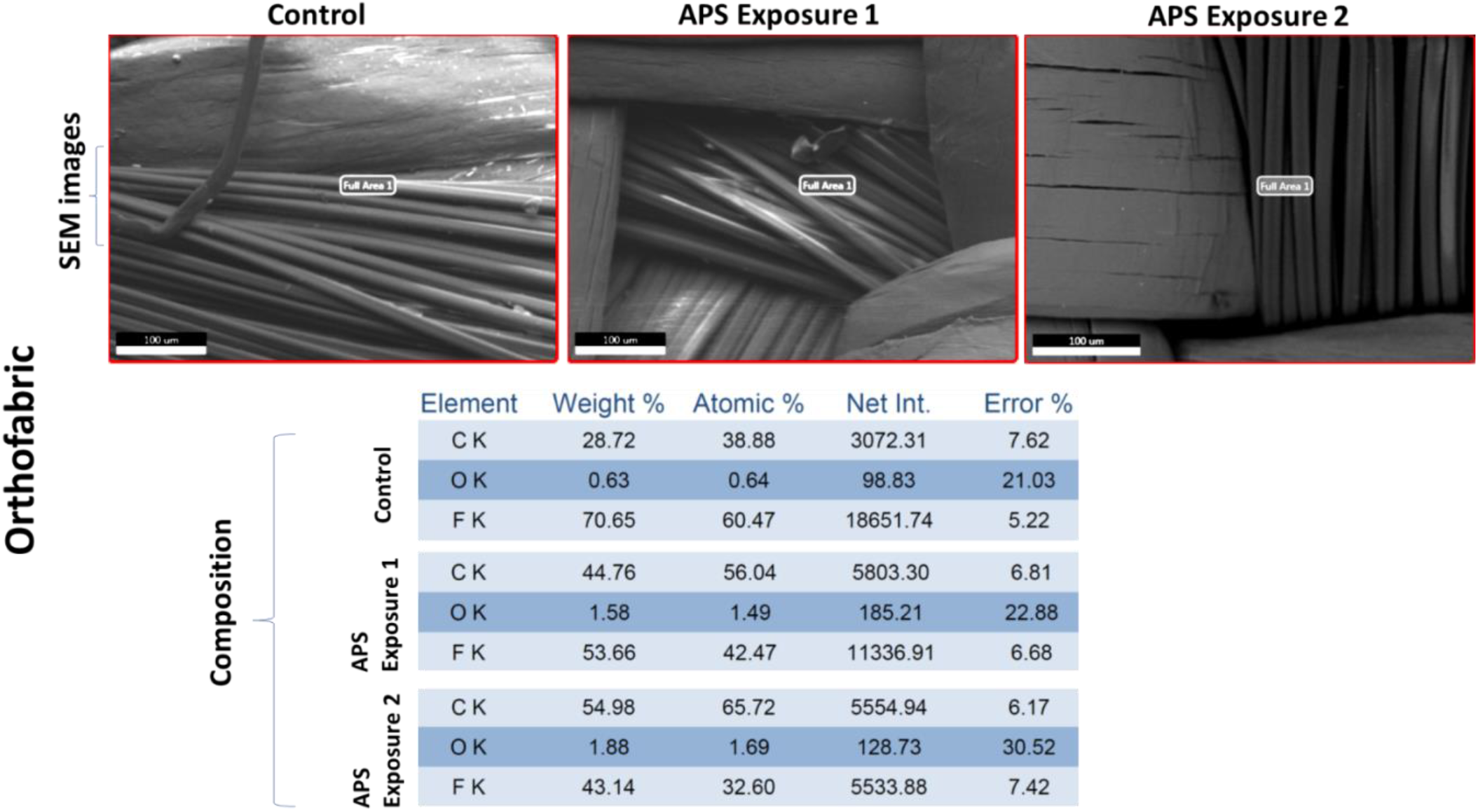
Material compatibility data: SEM analysis of Orthofabric with APS exposure 1: exposed to ozone CT value of 6789.02 ppm-min, and APS exposure 2: exposed to ozone CT value of 10323 ppm-min.

**Figure 18.**
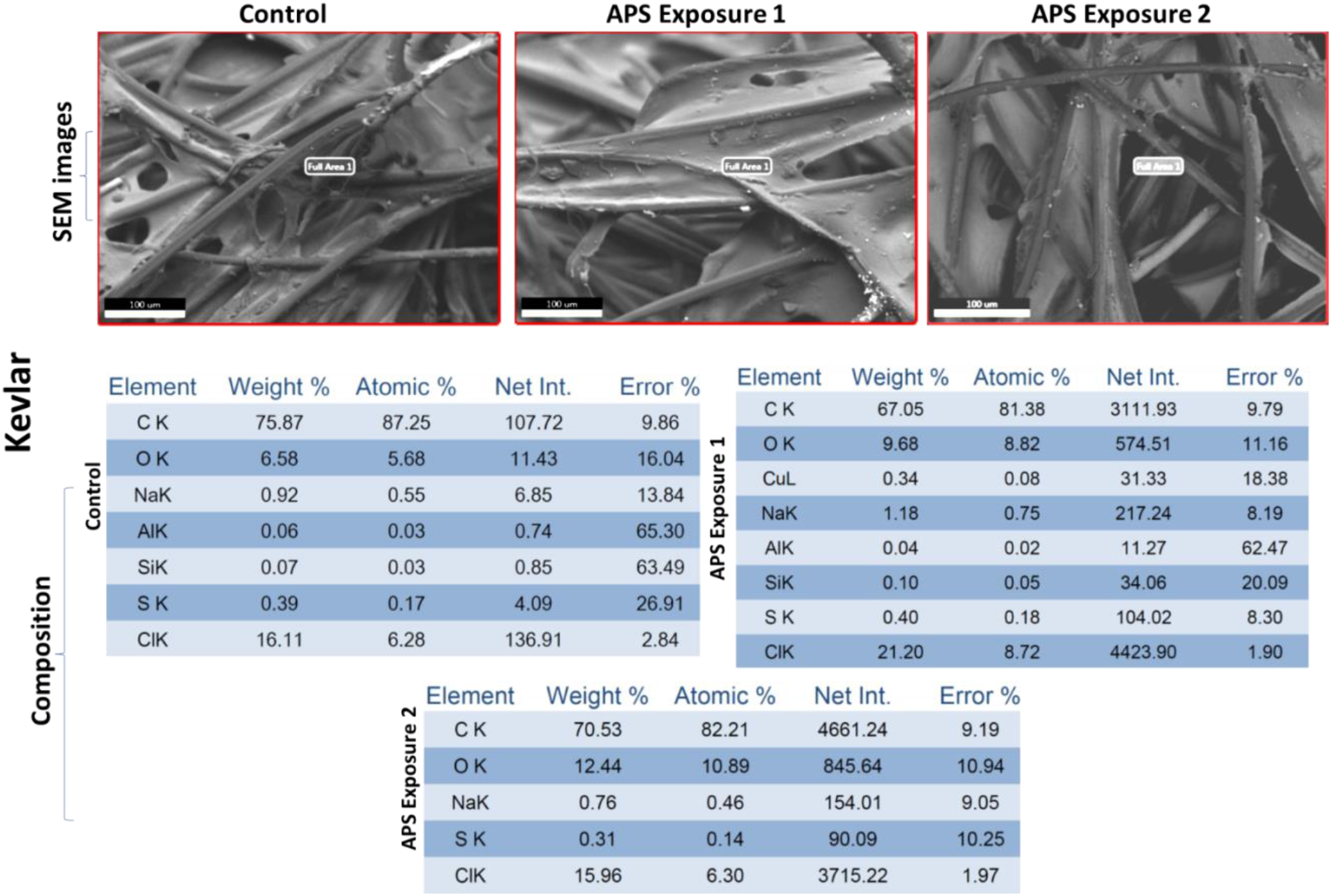
Material compatibility data: SEM analysis of Kevlar with APS exposure 1: exposed to ozone CT value of 6789.02 ppm-min, and APS exposure 2: exposed to ozone CT value of 10323 ppm-min.

## Conclusion

The effectiveness of the Active Plasma Sterilizer (APS) - a compact, sustainable and energy efficient sterilization system with inbuilt ozone generation and mixing capability was investigated for sterilization pertaining to planetary protection. Design of sterilization box of the first APS prototype (APS.V0) and integration of CPPRs in it for ozone generation and distribution was discussed. The CPPRs were integrated based on strategic selection and placement of SDBD reactor panel designs (Fan and Comb reactor configurations) to sustainably generate ozone using ambient air and for uniform distribution of the generated ozone within the sterilization box. The concept of achieving spatially distributed decontamination using a synergistic combination of SDBD ozone generation and flow actuation for distributing and mixing DBD generated ozone without using external mixing agents or mechanical moving parts was used. The APS prototype was evaluated by (a) testing its sterilization efficacy for coupons made of 4 material types (Aluminum, Polycarbonate, Kevlar, Orthofabric), contaminated with 4 to 5 logs of two test organisms: (*E.coli* and *B.subtilis*), and distributed at 11 points inside the APS, (b) determining the number of CPPRs required for complete killing of selected organisms within 30 minutes, (c) determining corresponding ozone CT (concentration x time) requirements and power requirements, (d) testing ozone penetration through selected fabric materials and (e) performing preliminary material compatibility tests through SEM analysis. Results show that the APS can achieve 4 to 5 log reductions of pathogenic bacteria (*E.coli* and *B.subtilis*) on Aluminum, Polycarbonate, Kevlar and Orthofabric, simultaneously at 11 points inside the chamber, within 30 minutes with 13.2 W total power consumption. Specifically, 4 to 5 log reductions of *E.coli* on all four materials was achieved within 20 minutes with ozone CT value of 6789.02 ppm-min; and 4 to 5 log reductions of *B.subtilis* on all four materials was achieved within 30 minutes with ozone CT value of 10323 ppm-min. Spatial distribution of the sterilization data at the central plane of the APS established that the APS can uniformly sterilize several points on a contaminated surface. Successful penetration of CPPR generated ozone for single and combined fabric layers was established with a maximum reduction of 16.17% in ozone concentrations through the layers without using an external agent or mechanical device to enhance penetration. Preliminary material compatibility tests with SEM analysis of 4 selected materials exposed to ozone CT values required for sterilization of both bacterial species in the APS showed no significant material damage. Thus, this study shows the potential of the APS as a sterilization technology for applications in planetary protection with advantages of uniform spatial decontamination, low processing temperatures, low exposure times, lightweight with no moving parts, ability to decontaminate porous surfaces and compatibility with relevant materials. Additionally, it is important to note that the DBD reactors has been successfully tested and found to regain performance even when water droplets are present on the work surface [30] making it a better candidate for a broad range of practical sterilization applications. Future research on the APS is needed for detailed material compatibility studies and further evaluation of the system in sterilizing other species of interest to planetary protection.

## Author Contributions

BC, and Subrata R contributed to conception of the study. BC, TR, SP and Subrata R contributed to development of the methodology. BC, TR, ENM, Sarthak R, and ML collected the data. BC, and Sarthak R made the figures. BC wrote the first draft of the manuscript. All authors contributed to manuscript revision.

## Acknowledgments

We acknowledge Dr. Lynn Torres of NASA and Dr. Alvin Smith from the Jet Propulsion Laboratory for their guidance in the project. We also acknowledge Nathan Rizza from Surfplasma for helping us in data collection.

